# PathX-CNN: An Enhanced Explainable Convolutional Neural Network for Survival Prediction and Pathway Analysis in Glioblastoma

**DOI:** 10.1101/2025.01.24.634827

**Authors:** Masrur Sobhan, Md Mezbahul Islam, Ananda Mohan Mondal

## Abstract

**Motivation:** Convolutional neural networks (CNNs) offer potential for analyzing non-grid structured data, such as biological array data, by converting it into image-like formats using principal component analysis (PCA) of pathway genes. However, PCA-derived principal components (PCs) from the entire dataset capture global variance but fail to extract sub-cohort (class-specific) variances. Consequently, CNNs trained on global PCs perform poorly in survival prediction of glioblastoma multiforme (GBM), and the corresponding explanation of CNN outcomes may not align with disease-relevant pathways.

**Results:** We present PathX-CNN, an explainable CNN framework that addresses these limitations by integrating multi-omics data through pathway-based images derived from sub-cohort-specific PCs. PathX-CNN outperformed existing pathway-based methods in predicting long-term survival (LTS) versus non-LTS in GBM. By leveraging SHAP (SHapley Additive exPlanations), a cooperative game theory-based explainable AI method, PathX-CNN identified biologically plausible pathways associated with GBM survival. Additionally, experiments on other cancer types demonstrated superior performance compared to traditional approaches. PathX-CNN demonstrates the potential of CNNs for multiomics integration, offering both improved prediction accuracy and pathway-specific insights into disease mechanisms.

**Code Availability:** https://github.com/codebysobhan/PathX-CNN

**Supplementary information:** Supplementary data are available at *Bioinformatics* online.

## 1 Introduction

Glioblastoma multiforme (GBM) remains the most aggressive and lethal form of primary brain tumor, with a median survival of approximately 12– 15 months despite advancements in surgical techniques, radiotherapy, and chemotherapeutic interventions [1], [2]. The heterogeneity of GBM, both at the inter-tumoral and intra-tumoral levels, significantly complicates treatment strategies and contributes to poor survival outcomes [3]. Accurate survival prediction is crucial for enabling personalized treatment strategies, optimizing therapeutic interventions, and improving overall clinical outcomes. However, the complex molecular mechanisms driving GBM progression and therapeutic resistance remain poorly understood, due to the complexity of its underlying biological pathways and the dynamic interplay of genetic, epigenetic, transcriptomic and environmental factors [4], [5].

The advent of multi-omics technologies, including genomics, transcriptomics, epigenomics, and proteomics, has provided unprecedented opportunities to dissect the molecular landscape of GBM and uncover key pathways associated with tumor progression and patient survival [6]. By integrating different layers of biological information, multi-omics data offer a more comprehensive understanding of GBM compared to single-omics approaches. However, despite significant progress, existing computational frameworks often fall short in effectively integrating these multiomics datasets, capturing non-linear relationships, and offering interpretable results for clinical translation [7]. There is a critical need for advanced computational models capable of leveraging multi-omics data to improve survival prediction and pathway analysis while providing transparent and interpretable outputs for clinical decision-making.

Current pathway-based computational methods for survival prediction and pathway analysis in glioblastoma (GBM) have made significant improvements, yet they are hindered by critical limitations that affect in clinical and translational research. PathCNN [8], an ideal example, attempted to integrate multi-omics data using a convolutional neural network (CNN) approach. However, it suffers from key methodological shortcomings. First, PathCNN employed principal component analysis (PCA) across all samples for dimensionality reduction and multi-omics integration. While PCA effectively reduces data dimensionality, applying it across all samples assumes global linear relationships and overlooks the heterogeneity of sub-cohorts or subtypes, which is crucial in understanding GBM’s diverse molecular profiles. Additionally, PathCNN evaluated predictive performance solely based on the area under the curve (AUC) score. While AUC provides an aggregate measure of classification performance, it is inherently limited in capturing nuances within imbalanced datasets, particularly in smaller sub-cohort with short-term survival. AUC tends to favor overall model performance while masking poor predictive power for underrepresented groups. This limitation raises concerns about the model’s clinical applicability, as survival predictions may not be reliable for minority cohorts or less prevalent survival patterns. Also, methods such as DeepKEGG [9] have improved pathway-based analysis by introducing hierarchical biological modules and self-attention mechanisms to better integrate sample-specific and pathway-level information. However, despite these improvements, DeepKEGG and similar models often prioritize overall performance metrics over sub-cohort-specific granularity, leaving room for biases that can obscure subgroup-level insights.

Addressing these limitations requires a new methodological framework that can: (1) preserve sub-cohort-specific heterogeneity during dimensionality reduction, (2) evaluate performance metrics beyond AUC, including sub-cohort-specific measures, and (3) ensure biological interpretability. This underscores the need for an enhanced explainable machine learning approach to address these persistent shortcomings in existing pathway-based survival prediction frameworks.

In diseases like glioblastoma (GBM), pathway-level analysis captures complex interactions and coordinated dysregulations [10], [11]. Integrating multi-omics data via pathway-level information reduces data dimensionality (the total number of pathway genes are ∼3,000 compared to ∼20,000 genes at the omics level) while preserving biologically meaningful patterns, improving both model interpretability and predictive accuracy. Furthermore, pathways often represent conserved mechanisms, enabling insights to be translated across multi-omics data [12]. By focusing on pathway-level features, machine learning models can better identify disease-driving mechanisms, enhance survival predictions, and provide actionable insights for GBM.

While machine learning models have shown good predictive performance (in term of AUC) in glioblastoma (GBM) survival analysis [8], a critical gap remains in their ability to predict and explain the smaller subgroup with short-term survival. The reason could be that the model like PathCNN rely on global dimensionality reduction techniques that obscure molecular signals specific to smaller subgroup with short-term survival. Traditional explainable AI (XAI) methods, such as Grad-CAM [13] as used in PathCNN, primarily focus on highlighting broad regions of importance without offering fine-grained, gene-level insights of pathway contributions. These approaches often fall short in tracing the exact contribution of specific features to the model’s predictions, limiting their utility in uncovering biological insights. In contrast, SHapley Additive exPlanations (SHAP) [14] have emerged as a more robust XAI framework, offering feature-level attributions [15], [16], that quantify each feature’s contribution to a model’s output in a mathematically grounded interpretable manner.

In this study, we propose PathX-CNN, an innovative explainable CNN framework that addresses the limitations of existing approaches by integrating multi-omics data through pathway-based images derived from sub-cohort-specific PCs and leveraging SHAP-based XAI.

Contributions of our work.

1. Improved pathway-image generation: Class-specific PCs are used to generate pathway images instead of global PCs, which is innovative and a major contribution of this work.
2. Integration of SHAP-based AI with CNN-based survival predictions, which provides class-specific biomarkers and pathway dependencies.
3. A survival prediction framework with improved performance. Beyond GBM, we demonstrate the model’s broader applicability across other disease types (KIRC, LGG, LUAD, and LUSC), showcasing its robustness and versatility in diverse multi-omics datasets.

The remainder of this paper is organized as follows: Materials and Methods section describes the datasets used with their summary statistics, pathway embedding process, the architecture of the PathX-CNN model, and the explainability approach leveraging SHAP. The Results section presents the predictive performance of PathX-CNN across different evaluation metrics and provides insights from pathway analysis. We also compared our approach with the benchmark methods. This section also highlights key pathways associated with glioblastoma (GBM) survival. In the Discussion, we explained the broader implications of our findings for multi-omics data integration, and the potential impact on GBM research and clinical practice. Finally, the Conclusion section summarizes the key contributions of this study, emphasizing the improved accuracy and interpretability of PathX-CNN, and outlines potential future research directions.

## 2 Materials and Methods

### 2.1 Data Collection

The datasets utilized in this study were sourced from cBioPortal [17].Three types of omics data were incorporated to ensure a comprehensive analysis of glioblastoma (GBM): Copy Number Variation (CNV), Gene Expression (EXP), and DNA Methylation (MET) from both hm27 and hm450 platforms.

Pathway information, along with the associated genes for each pathway, was downloaded from MSIGDB KEGG_LEGACY [18] database consisting of 186 pathways. After excluding disease-specific pathways, a total of 146 pathways were selected for analysis. They were excluded to focus on fundamental biological processes and pathways common across various cellular contexts and organisms. Excluding these pathways helps avoid potential bias in the analysis, as they are tailored to specific diseases and may not accurately represent general biological processes.

### 2.2 Data Preparation

To harmonize the multi-omics data, preprocessing was performed to identify common samples and genes across the three omics data types.

The raw datasets contained varying number of samples and features-- CNV: 577 samples with 24,776 features, EXP: 528 samples with 12,042 features, MET-HM27: 285 samples with 11,807 features, and MET-HM450: 153 samples with 11,807 features. After combining the two methylation platforms, MET data contains 433 samples with 11,805 features. Finally, the processed dataset consists of 347 common samples with 8,044 common features across all three omics types. Long-term survival (LTS) was defined as survival > 2 years after diagnosis, while non-LTS was defined as survival ≤ 2 years. Individuals with a last follow-up of ≤ 2 years were excluded from further analysis. This thresholding resulted in 55 LTS, 234 non-LTS, and 58 excluded samples.

### 2.3 Study Flow Diagram

Figure 1 shows the flow diagram of the proposed framework, PathX-CNN, an enhanced explainable method for survival analysis by integrating multi-omics data via the embedding of omics information through pathways. In this study, we integrated three types of omics data: copy number variation (CNV), mRNA expression (EXP), and DNA methylation (MET). The integration process involved performing Principal Component Analysis (PCA) on each pathway genes separately for each omics type and for each class cohort (LTS and Non-LTS). In other words, class-specific PCA was conducted for each omics types. The samples, embedded using principal components (PCs), were the input to convolutional neural network (CNN) model. The trained CNN model, along with the Pathway-PC images, was subsequently fed into the explainable AI tool Gradient-SHAP to identify patient-specific significant pathways contributing in distinguishing LTS from non-LTS group. SHAP assigns significance scores to each of the PCs. The top-ranking PCs based on SHAP score are deemed significant, and the associated omics and pathways are identified as significant omics and pathways. Finally, we determined the significant genes within these significant pathways by Cox Proportional Hazards (CoxPH) model.

**Fig. 1.**
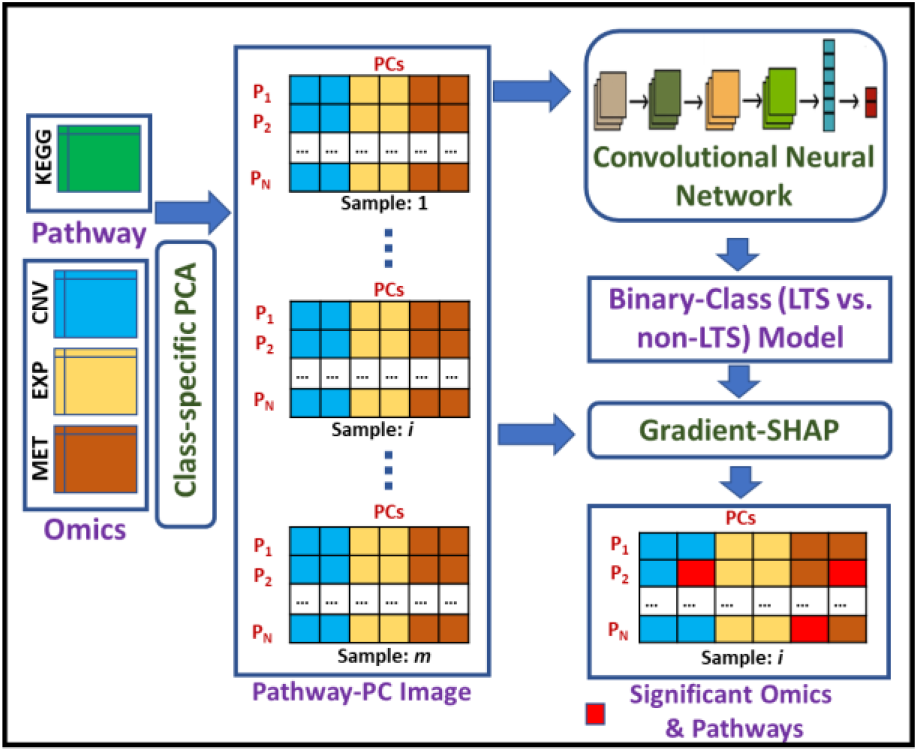
Overall Pipeline of PathX-CNN. CNV: Copy Number Variation, EXP: mRNA Expression, MET: DNA Methylation, PCA: Principal Component Analysis, PC: Principal Component, P_i_: Pathway.

### 2.4 Pathway-PC Image Generation

Figure 2 illustrates the steps to generate the Pathway-PC images.

**Fig. 2.**
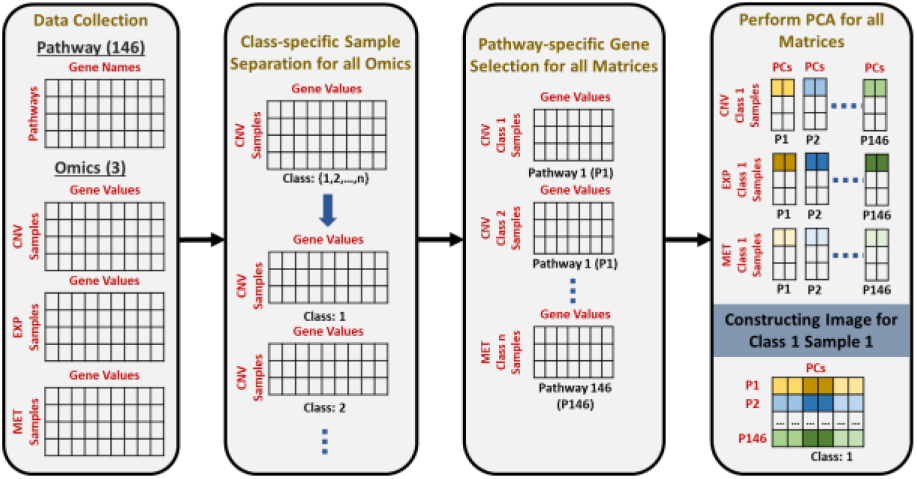
Generation of Pathway-PC images as the input to CNN Model.

***Data Collection*** box shows the pictorial representation of four different datasets used for analysis, including (i) 146 pathways with corresponding gene names, (ii) samples with CNV values for genes, (iii) samples with EXP values for genes, and (iv) samples with MET values for genes.

#### Class-specific Sample Separation for Each Omics

The objective of this step is to isolate samples for each class and each omics data type. For an n-class problem with *t* omics type, we need to generate *n* × *t* datasets. The present study is a 2-class problem (LTS vs. non-LTS) with 3 omics data types (CNV, EXP, and MET), which will result in 6 2-D matrices.

#### Isolating Pathway-specific Gene Values

This step isolates gene values for each pathway from each of the 2-D matrices derived in previous steps, which will result in 146 pathway-specific 2-D matrices for each class and each omics type. Altogether, it will generate (2 × 3 × 146 =) 876 2-D matrices.

#### Perform PCA for all Matrices

Principal Component Analysis (PCA) was applied on each of the 876 matrices derived in the previous step. The goal of this step is to integrate the multi-omics data by bringing it to the same level (for example, 2 PCs from each omics and each pathway), allowing for a unified analysis across different data types. By focusing on the top PC components, we aimed to capture the most significant information from each pathway, thereby reducing the dimensionality of the features and addressing the curse of dimensionality. Since the dimensionality reduction or the embeddings was done on each class separately, it preserves the class-specific biological signals.

#### Constructing Pathway-PC Matrix/Image

In general, Pathway-PC matrix/image for a sample can be represented as (*number of pathways*× (*number of omics*× *number of PCs*)). The bottom part of box 4 in Figure 2 shows the construction of Pathway-PC matrix/image (146 × 6) for the sample 1 of class 1 (class LTS) using top 2 PCs from three omics.

### 2.5 Architecture of Convolutional Neural Network

CNN model in PathX-CNN framework consists of 4 convolutional layers as shown in *Supplementary* Figure S1. Each of the convolutional layers are followed by batch normalization, activation, and MaxPooling layers. The final convolutional outputs are flattened into a one-dimensional representation, enabling the transition from spatial feature extraction to fully connected layer. Dense layer is applied to capture global pathway-level interactions, accompanied by batch normalization and non-linear activation functions to ensure robust learning and generalization. The final output layer produces class probabilities using a softmax activation function. The model is compiled with the Adam optimizer for efficient gradient-based learning. Accuracy is used as the primary evaluation metric to monitor the model’s performance. The model was trained for 100 epochs with a batch size of 32 and a validation split of 20%. This setup enabled effective learning while monitoring validation performance to prevent overfitting.

To rigorously evaluate the classification performance, we adopted a stratified 5-fold cross-validation strategy. We conducted hyperparameter tuning using Bayesian Optimization technique to optimize the architecture and training process. Details of hyperparameter tuning is provided in *Supplementary* Table S1.

### 2.6 Evaluation Metrics

#### Accuracy

Accuracy is the ratio of correctly predicted instances to the total number of instances. It is calculated as accuracy= (TP + TN) / (TP + TN + FP + FN), where TP, TN, FP, and FN are true positives, true negatives, false positives, and false negatives, respectively. While accuracy is a simple and widely used metric, it may be misleading for imbalanced datasets, as it does not differentiate between the types of errors.

#### Precision

Precision measures the proportion of true positive predictions out of all positive predictions made by the model. It is calculated as: precision= (TP) / (TP + FP).

#### Recall

Recall, also known as sensitivity or true positive rate, calculates the proportion of actual positive instances correctly identified by the model. It is calculated as: recall= (TP) / (TP + FN).

#### F1-score

The F1-score is the harmonic mean of precision and recall, providing a single metric that balances both. It is defined as: f1-score = 2 × (precision × recall) / (precision + recall)

#### AUC-ROC score

The AUC-ROC score quantifies the area under the Receiver Operating Characteristic (ROC) curve, which plots the true positive rate (recall) against the false positive rate (1 - specificity) at various thresholds. A higher AUC-ROC score indicates better model performance in distinguishing between positive and negative classes, with a perfect classifier scoring 1.0.

#### Confusion Matrix

The confusion matrix is a tabular representation of a model’s performance, showing the counts of TP, TN, FP, and FN. It provides a detailed breakdown of classification results, enabling a deeper understanding of model errors and successes.

### 2.7 Feature Interpretation using Gradient SHAP

SHAP is a powerful explainable AI (XAI) tool based on game theory that provides insights into how machine learning models make decisions. It assigns a score to each feature for every sample, indicating the extent to which each feature influences the model’s prediction. The fundamental concept behind SHAP is to assess the marginal contribution of a feature by comparing the model’s output with and without the feature, across various combinations of features called coalition sets. This process ensures that each feature’s contribution is evaluated in the context of all possible feature subsets.

In our study, we employed Gradient SHAP for feature explanation since our model utilizes a Convolutional Neural Network (CNN)-based classifier. Gradient SHAP [19] leverages the gradients of the model’s output with respect to the input features, making it particularly suited for differentiable models like CNNs. It incorporates multiple baselines by perturbing the input data with noise. This reduces sensitivity to baseline selection and improves the robustness of the explanations. The Gradient SHAP algorithm calculates the attribution of a feature *x*_*i*_ by averaging the gradients of the model’s output along paths from multiple baselines *x’* to the actual input *x*_*i*_. The SHAP value for feature *x*_*i*_ is defined as:

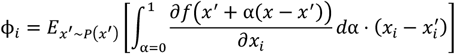

where *x’* is a baseline input sampled from a distribution *P*(*x*^′^), *x* is the actual input being explained, α is a scaling factor that interpolate between the baseline (*x*^′^ + α = 0) and the input (*x*^′^ + α = *x*), *f*(*x*) is the prediction of the model for input *x* and 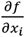 is the gradient of the model’s prediction with respect to feature *x*_*i*_. The formula ensures that the contribution of each feature is assessed over all possible paths between the baseline and the input, capturing the feature’s influence comprehensively. Gradient SHAP generates these paths stochastically by adding noise to the input, improving the explanation’s robustness.

#### Significant Omics and Pathways

In this study, based on the global interpretation of SHAP, top 10 features (PCs in this study) for each class are identified. In our case, the features are the PCs in different omics derived from pathways. We can roll back to significant omics and pathways from the significant PCs as shown in the “Significant Omics and Pathways” component of the flow diagram. Thus, the proposed approach, PathX-CNN, can find both significant omics and pathways in differentiating LTS from non-LTS patients.

### 2.8 Cox Proportional Hazards Model to identify Significant Genes from Significant Pathways

The CoxPH model estimates the relationship between a set of covariates and the hazard rate, which is the instantaneous probability of an event (e.g., death) occurring at a given time. The hazard function *h*(*t*|***X***) in the CoxPH model is given by:

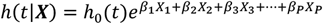

where *h*(*t*|***X***) is hazard rate for an individual with covariates (here, genes) *X = (X*_*1*_, *X*_*2*_, *X*_*3*_,*…*., *X*_*p*_) from significant pathways, *h*_0_(*t*) is the baseline hazard rate shared across all the individuals, and *β* represents coefficients associated with each covariate. To identify whether a covariate (e.g., a specific gene) significantly impacts survival a hypothesis test is performed where:

Null hypothesis (H_0_) is: the covariate has no effect on survival (*i. e*., *β* = 0) and alternative hypothesis (H_a_) is: the covariate affects survival (*i. e*., *β* ≠ 0). The significance is assessed using a p-value derived from the Wald test. The Wald statistic is calculated as:

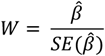

where 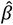 is the estimated coefficient for the covariate (a gene) and 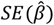 is the standard error of the estimated coefficient. The p-value is then derived from the Wald statistic using a standard normal distribution. If the p-value is below a chosen threshold, the covariate is considered statistically significant. In this study, p-value is chosen < 0.01. Finally, the hazard ratio was evaluated for a significant covariate (gene) using the expression, *e*^*β*^, where *β* is the coefficient of the significant covariate.

### 2.9 Dataset for Validating the Pipeline

We utilized publicly available multi-omics datasets from The Cancer Genome Atlas (TCGA) to validate the generalizability of the proposed approach. We downloaded datasets for kidney renal clear cell carcinoma (KIRC), low-grade glioma (LGG), lung adenocarcinoma (LUAD) and lung squamous cell carcinoma (LUSC) from Xena Browser [20]. These datasets include genomic (gene level copy number & gene-level non-silent mutation), transcriptomic (RNA sequencing gene expression), and epigenomic (DNA methylation: 27k & 450k) data. Note that for validating the pipeline we are using one additional omics data (mutations). The summary of the processed data is shown in Table 1.

**Table 1.** Brief data description for various cancer types.

| Can-<br>cer | Features |  |  |  | Total Samples<br>(LTS : non-LTS) |
| --- | --- | --- | --- | --- | --- |
|  | CNV | EXP | MET | MUT |  |
| KIRC | 24,776 | 20,530 | 25,978 | 40,543 | 249 (188 : 61) |
| LGG | 24,776 | 20,530 | 485,577 | 40,543 | 228 (153 : 75) |
| LUAD | 24,776 | 20,530 | 25,978 | 40,543 | 244 (121 : 123) |
| LUSC | 24,776 | 20,530 | 25,978 | 40,543 | 274 (134 : 140) |

### 2.10 Comparison with Benchmark Methods

The predictive performance of PathX-CNN was evaluated against several established machine learning models, including Logistic Regression, Naive Bayes Classifier, PathCNN, and DeepKEGG. To ensure a comprehensive comparison, multiple evaluation metrics were used, including accuracy, precision, recall, F1-score, AUC score, and confusion matrix. These metrics provided a well-rounded assessment of both the predictive accuracy and the robustness of each classifier.

## 3 Results

### 3.1 Optimal Number of Principal Components (PCs)

The selection of the optimal number of principal components (PCs) was done using the criterion of achieving the highest AUC score while minimizing the number of PCs to reduce computational complexity. From the observed trend shown in Figure 3, the AUC score increases significantly from 1 PC to 2 PCs, reaching its peak at 2 PCs with an AUC score close to 99%. Beyond this point, adding more PCs results in a decline in performance. Therefore, the optimal number of PCs is 2, which was used for subsequent analyses.

**Fig. 3.**
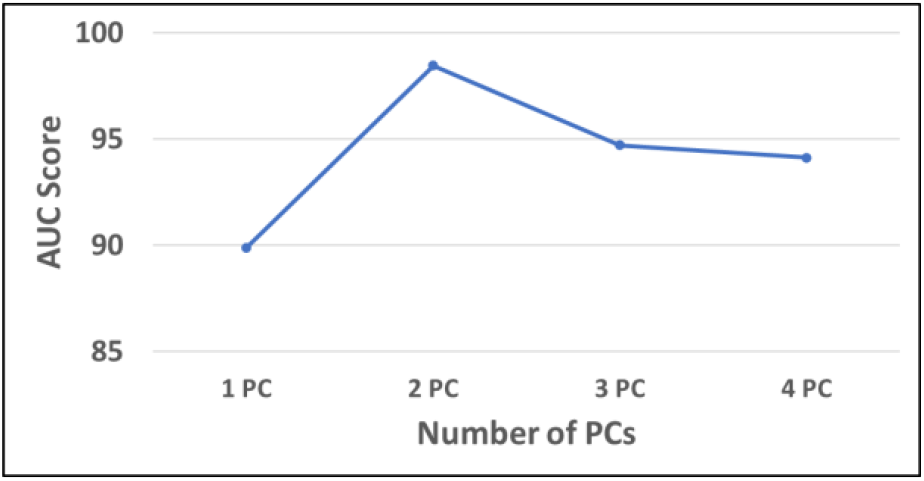
Performance Comparison Across Different Numbers of Principal Components (PCs). The AUC scores were evaluated across one to four principal components (PCs) to identify the optimal number for model performance. The highest AUC score was achieved using 2 PCs.

### 3.2 Performance Evaluation with Benchmark Methods

We evaluate the performance of PathX-CNN against four benchmark methods: Deep-KEGG, Logistic Regression (LR), Naive Bayes (NB), and PathCNN across five cancer types: GBM (Glioblastoma Multiforme), KIRC (Kidney Renal Clear Cell Carcinoma), LGG (Lower Grade Glioma), LUAD (Lung Adenocarcinoma), and LUSC (Lung Squamous Cell Carcinoma). The evaluation focuses on five key metrics: Accuracy (Acc), Precision (Pre), Recall (Rec), F1 Score (F1), and AUC Score (AUC) shown in Table 2.

**Table 2.** Comparison of predictive performance with benchmark methods in terms of accuracy (Acc), precision (Pre), recall (Rec), f1-score (F1), and area under the curve (AUC).

| Dis-<br>ease | Met-<br>rics | Deep-KEGG | LR | NB | PathCNN | PathX-CNN |
| --- | --- | --- | --- | --- | --- | --- |
| GBM | Acc | 75.77% | 80.96% | 71.97% | 80.88% | **95.5%** |
|  | Pre | 84.74% | 80.96% | 86.60% | 81.64% | **96.62%** |
|  | Rec | 85.47% | **100%** | 77.35% | **98.49%** | 97.86% |
|  | F1 | 85.10% | 89.48% | 81.71% | 89.28% | **97.23%** |
|  | AUC | 63.76% | 26.87% | 65.83% | 74.39% | **98.45%** |
| KIRC | Acc | 73.89% | 75.50% | 86.75% | 76.31% | **89.56%** |
|  | Pre | 47.05% | NA | **88.89%** | 60.00% | 87.23% |
|  | Rec | 52.45% | 0% | 52.45% | 9.84% | **67.21%** |
|  | F1 | 49.61% | NA | 65.98% | 16.90% | **75.93%** |
|  | AUC | 72.13% | 16.36% | 76.55% | 67.05% | **82.13%** |
| LGG | Acc | 80.70% | 67.10% | 80.70% | 86.84% | **94.74%** |
|  | Pre | 70.67% | NA | 75.40% | 86.89% | **97.01%** |
|  | Rec | 70.67% | 0% | 61.33% | 70.67% | **86.67%** |
|  | F1 | 70.67% | NA | 67.65% | 77.94% | **91.55%** |
|  | AUC | 84.29% | 21.61% | 75.72% | 84.65% | **92.68%** |
| LUAD | Acc | 56.96% | 36.49% | 36.08% | 60.66% | **91.80%** |
|  | Pre | 57.89% | 40.90% | 37.59% | 61.95% | **96.40%** |
|  | Rec | 53.65% | 58.53% | 40.65% | 56.91% | **86.99%** |
|  | F1 | 55.69% | 48.16% | 39.06% | 59.32% | **91.45%** |
|  | AUC | 58.67% | 24.38% | 29.78% | 65.22% | **91.80%** |
| LUSC | Acc | 54.75% | 39.21% | 54.56% | 52.55% | **91.24%** |
|  | Pre | 55.47% | 40.86% | 47.82% | 53.38% | **90.28%** |
|  | Rec | 57.85% | 89.95% | 52.63% | 56.43% | **92.86%** |
|  | F1 | 56.64% | 56.20% | 50.11% | 54.86% | **91.55%** |
|  | AUC | 52.50% | 30.07% | 50.21% | 53.56% | **91.22%** |

In GBM, PathX-CNN outperforms all other models in accuracy (95.5%), precision (96.62%), F1 Score (97.23%), and AUC score (98.45%) in distinguishing non-LTS (positive class) from LTS (negative class), indicating a strong balance between identifying true positives and true negatives while minimizing false positives and false negatives, evidenced from confusion matrix (*Supplementary Figure S2*). PathCNN follows closely with 98.49% recall, meaning it correctly identifies nearly all non-LTS patients, but its precision (81.64%) is lower, suggesting a higher number of false positives compared to PathX-CNN. Among traditional machine learning methods, Logistic Regression (LR) achieves 100% recall, meaning it classifies all patients (non-LTS + LTS) as positive (non-LTS), but this comes at the cost of a lower precision (80.96%), leading to a high number of false positives. Naïve Bayes (NB) shows moderate performance with 86.60% precision and 77.35% recall, resulting in an F1 Score of 81.71%. Deep-KEGG demonstrates comparable results to NB, but with slightly better recall (85.47%) and a lower precision (84.74%), leading to an F1 Score of 85.10% which also indicates a moderate performance. The overall trend indicates that PathX-CNN is the most balanced model, achieving the highest F1 (97.23%) and AUC (98.45%) scores, which confirms that it maintains both high precision and recall. In contrast, PathCNN tends to favor recall over precision, while Logistic Regression completely prioritizes recall, making it less reliable for precise classification. Naïve Bayes and DeepKEGG struggle to balance sensitivity and specificity, making them less effective as prediction models. Based on all evaluation metrics, the proposed method PathX-CNN outperforms all the benchmark models in predicting both non-LTS and LTS GBM patients.

#### Robustness of PathX-CNN

To check the robustness of the proposed PathX-CNN, the whole experiment was repeated with four other cancers, including KIRC, LGG, LUAD, and LUSC.

PathX-CNN outperforms all classifiers across all metrics, except for precision in KIRC, where Naïve Bayes (88.89%) slightly surpasses PathX-CNN’s precision (87.23%). But Naïve Bayes suffers with a low recall (52.45%) compared to PathX-CNN (67.21%). For KIRC, PathX-CNN once again outperforms other models maintaining a good balance between precision and recall. PathCNN struggles with recall (9.84%), highlighting its poor ability to avoid false negatives. Logistic Regression (LR) did not yield precision and F1 Score values because both true positive (TP) and false positive (FP) counts were zero, leading to an undefined precision metric. In LGG, LUAD, and LUSC, PathX-CNN significantly outperformed all other classifiers, demonstrating the robustness and effectiveness of this framework in accurately distinguishing non-LTS and LTS patients.

### 3.3 Identifying Significant Pathways for GBM

In this study, we leveraged the Gradient SHAP explainer to identify significant pathways contributing to survival prediction (non-LTS and LTS) in the PathX-CNN framework as this classifier performed better compared to other classifiers. The Gradient SHAP method enabled us to compute SHAP values by approximating feature importance through the gradients of the PathX-CNN model output with respect to input features. These SHAP values provided a detailed interpretation of the model’s predictions, highlighting the pathways that played a critical role in differentiating between long-term survival (LTS) and non-long-term survival (non-LTS) groups.

For each class, we computed the global scores by averaging the SHAP values across all samples within that class. This approach ensured that the identified pathways captured class-specific information, reflecting the most influential biological processes contributing to the predictions for each survival group. Employing class-specific global interpretations of SHAP,we identified top 10 significant pathways that are biologically meaningful within the context of non-LTS (*Supplementary Figure S3*) and LTS (*Supplementary Figure S4*) patients, Among these two sets of top 10 pathways, six pathways were found to be common, as illustrated in the Venn diagram of Figure 4a. This overlap is expected, as both classes originate from the same disease, and these six pathways likely contain the most informative and discriminative features that contribute to distinguishing between the LTS and non-LTS groups. The overlap between the two groups indicates shared biological processes relevant to both survival classes. Notably, the MAPK Signaling Pathway emerged twice as significant, and the both PC values are from the mRNA expression profile. Additionally, four unique pathways are exclusive to each group, emphasizing distinct biological characteristics that differentiate the non-LTS and LTS groups. Figure 4b presents a table listing the names of the pathways categorized as common and unique. The “Common Pathways” column contains the overlapping pathways, while the “Non-LTS Pathways” and “LTS Pathways” columns show the four unique pathways for each group.

**Fig. 4.**
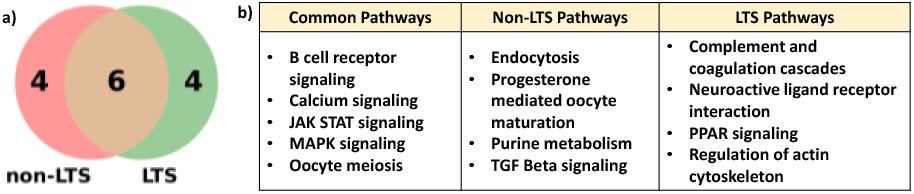
Comparative Analysis of Significant Pathways for Non-LTS and LTS Groups. (a) Venn diagram highlighting the shared and unique pathways among the top 10 significant pathways. (b) Pathway names, categorized by the common pathways and the pathways unique to non-LTS and LTS groups.

### 3.4 Identifying Significant Genes from Significant Pathways

Figure 5 presents a forest plot showing the significant genes within the oocyte meiosis pathway that are shared between Long-Term Survivors (LTS) and Non-Long-Term Survivors (non-LTS) of Glioblastoma Multiforme (GBM). The analysis was performed using the Cox Proportional Hazards (CoxPH) model, which estimates the association between gene expression levels and patient survival. A statistical threshold of p-value < 0.01 was applied to determine significance. The y-axis lists the genes identified as significant within the oocyte meiosis pathway in GBM based on p-values. The x-axis represents the Hazard Ratio (HR), which quantifies the impact of gene expression on survival. HR > 1 indicates that higher expression of the gene is associated with a higher risk of death (worse prognosis). HR < 1 suggests that higher expression of the gene is associated with a lower risk of death (better prognosis). The black horizontal bars represent the 95% Confidence Intervals (CIs), which indicate the range of HR values within which the true effect likely lies. From the figure, it is clear that PPP2R1B (HR = 3.28) has the strongest predictor of poor survival, with its higher expression correlating with a significantly increased risk of death. NAPC10 (HR = 1.98), IGF1 (HR = 1.85), and IGF1R (HR = 1.52) also show a positive hazard ratio, suggesting they contribute to GBM progression and poor prognosis. PRKACB (HR = 0.586), PPP2R5C (HR = 0.388), and MAPK3 (HR = 0.143) are associated with improved survival, suggesting their higher expression is protective against tumor progression. Forest plots of other common significant pathways and the names of the genes are provided in *Supplementary Figures S7 through S11*.

**Fig. 5.**
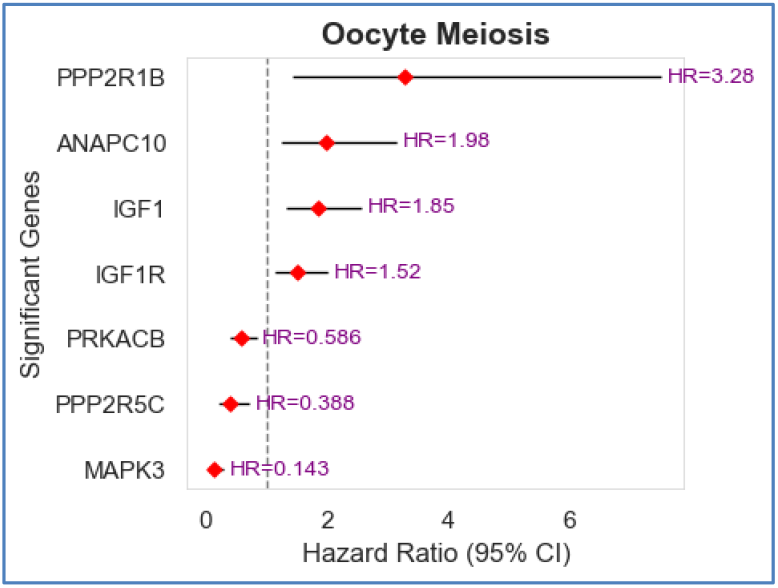
Hazard ratios and confidence interval of significant genes in the oocyte meiosis pathway associated with survival in GBM.

## 4 Discussion

We hypothesize that pathway-based multi-omics integration via the class-specific embedding of pathway information through PCA will perform better than the integration via the whole cohort-specific embedding. To test this hypothesis, we developed a novel framework called PathX-CNN to analyze the survival of GBM by creating two classes of LTS and non-LTS. Our approach performed way better than the whole cohort-specific embedding used in the state-of-the-art approach, PathCNN. The significance of class-specific pathway-based embeddings lies in their ability to preserve biologically relevant features across different omics types, enabling the model to make more accurate predictions. Our approach, PathX-CNN showed a great improvement across all the evaluation scores because of both class-specific PCA and the architecture of CNN.

The integration of SHAP-based interpretability allows for identifying significant pathways that contribute to model predictions. This interpretability is crucial for understanding the underlying biology of survival outcomes and can guide further experimental validation. In this study, we utilized the Gradient SHAP explainer to identify significant pathways that contribute to survival prediction within the PathX-CNN framework. Gradient SHAP approximated SHAP values that provided feature importance scores for each pathway. These scores enabled us to interpret the predictions of the PathX-CNN model effectively, identifying pathways that differentiate between long-term survival (LTS) and non-long-term survival (non-LTS) groups in GBM cancer. Global interpretation score was used in identifying top 10 significant pathways for both the LTS and non-LTS groups. Among these pathways, four unique pathways for each survival group, emphasizing class-specific biological characteristics that differentiate the LTS and non-LTS groups. These distinct pathways likely represent unique survival mechanisms or molecular changes that are critical to each group. The analysis identified six pathways were shared between the two classes, which are-B cell receptor signaling, Calcium signaling, JAK STAT signaling, MAPK signaling, and Oocyte meiosis pathways, indicating the presence of common biological processes that are critical to both survival groups. These overlapping pathways highlight core mechanisms that contribute to cancer progression and survival outcomes in GBM. Our findings in detecting significant pathways from PathX-CNN framework are validated by the existing literature reviews. Hou et. al. found that neutralizing TGFβ-mediated immunosuppression led to increased B cell infiltration and activation within the tumor microenvironment [21]. This B cell response was associated with improved therapeutic outcomes, suggesting that B cell receptor signaling plays a crucial role in modulating immune responses during PD-1 blockade treatment in GBM. Calcium signaling plays a pivotal role in glioblastoma (GBM) progression by influencing various cellular processes [22]. Alterations in calcium homeostasis can affect tumor cell proliferation, migration, and resistance to apoptosis, thereby contributing to the aggressive nature of GBM. The JAK/STAT3 signaling pathway plays a critical role in the pathogenesis and progression of GBM [23]. STAT3 expression is significantly elevated in GBM tissues compared to normal brain cells, and its abnormal activation contributes to an immunosuppressive tumor microenvironment in GBM. The MAPK pathway is frequently altered in glial tumors. In glioma cells, p38 MAPK activation promotes tumor invasion and metastasis, showing a positive correlation with tumor grade [24]. It is regarded as a potential oncogenic factor involved in brain tumorigenesis and chemotherapy resistance. In GBM, MAPK hyperactivation drives key processes like migration, proliferation, and survival by regulating cell proliferation and CREB activation, a cyclin-D1 regulator [25]. The Oocyte Meiosis pathway has been implicated in glioblastoma (GBM) through bioinformatics analyses identifying its involvement in the disease’s progression. Specifically, genes such as PRKCB have been found to be expressed in pathways including oocyte meiosis, suggesting a potential role in glioma development [26]. These findings suggest that targeting these pathways could influence tumor progression and survival outcomes in GBM.

Significant non-LTS pathways identified in our analysis have been validated by existing literature, further emphasizing their biological relevance in GBM progression and survival outcomes. Endocytosis plays a critical role in receptor recycling and signal transduction, which are essential for tumor growth and survival, as highlighted in a study that explores its dysregulation in GBM and its impact on cellular signaling networks [27]. Similarly, the progesterone-mediated oocyte maturation pathway, although primarily associated with reproductive processes, has been implicated in GBM through aberrant expression of pathway components that influence tumor behavior [28]. Purine metabolism, a critical pathway for nucleotide biosynthesis, supports the rapid proliferation of cancer cells, with research demonstrating its role in sustaining tumor growth and identifying it as a potential therapeutic target in GBM [29]. Lastly, the TGF-β signaling pathway has been extensively studied in GBM for its contributions to tumor invasiveness, immunosuppression, and maintenance of a tumorigenic microenvironment, making it a promising focus for therapeutic interventions [30].

PathX-CNN framework also identified significant pathways in the long-term survival (LTS) group of GBM patients. Complement and coagulation cascades have been implicated in immune regulation and tumor progression, with research showing their dual role in both enhancing and suppressing tumor growth, depending on the tumor microenvironment [31]. The neuroactive ligand-receptor interaction pathway, crucial for neurotransmitter signaling, has been associated with GBM progression and neural invasion, playing a role in the tumor’s ability to interact with and influence the brain’s normal neuronal processes [32]. PPAR signaling has emerged as a key metabolic regulator, with recent studies suggesting its potential to inhibit GBM cell proliferation and induce differentiation, offering a promising target for therapeutic interventions [33]. Lastly, the regulation of actin cytoskeleton pathway plays a fundamental role in cell migration, invasion, and metastasis, with its dysregulation being linked to enhanced invasive behavior in GBM cells [34].

The identification of significant genes using CoxPH model within critical pathways is essential for understanding disease mechanisms in glioblastoma (GBM). This process offers valuable insights into the molecular and biological factors that drive tumor progression, ultimately enabling the development of targeted therapeutic strategies. Significant genes act as key regulators within pathways that control essential cellular functions such as proliferation, apoptosis, migration, and immune evasion. Understanding how these genes interact within their pathways provides a deeper understanding of the processes that sustain tumor aggressiveness and drive GBM progression. Targeting significant genes also opens new therapeutic avenues. These genes can serve as potential drug targets, allowing therapies to disrupt key signaling pathways that support tumor survival and proliferation. Identifying significant genes within critical pathways is instrumental in advancing our understanding of GBM. These genes reveal key molecular drivers of tumor progression, act as biomarkers for survival prediction, and provide promising therapeutic target.

## 5 Conclusion

In this study, we introduced PathX-CNN, a pathway-based deep learning framework that integrates multi-omics data to address the complex challenge of survival prediction across glioblastoma multiforme (GBM), kidney renal clear cell carcinoma (KIRC), low-grade gliomas (LGG), lung adenocarcinoma (LUAD), and lung squamous cell carcinoma (LUSC). By employing class-specific Principal Component Analysis (PCA), PathX-CNN generated pathway-level embeddings that capture critical genomic and epigenomic patterns specific to distinct survival groups. This innovative integration strategy, coupled with explainable AI tool, SHAP, enables a more explainable analysis of biological pathways. The class-specific embedding not only enhances predictive accuracy but also reveals molecular insights that are essential for understanding cancer survival mechanisms. The superiority of PathX-CNN over traditional machine learning and deep learning models-such as PathCNN and DeepKEGG - was evident across all performance metrics, particularly in its ability to handle imbalanced datasets with precision and reliability. By leveraging SHAP-based interpretations, PathX-CNN uncovered significant pathways, including MAPK signaling, calcium signaling, B cell receptor signaling, and JAK/STAT signaling, elucidating their roles in tumor progression in GBM. The identification of shared and unique pathways across long-term survival (LTS) and non-long-term survival (non-LTS) groups highlights the framework’s ability to disentangle overlapping core processes from distinct molecular mechanisms driving survival differences. Furthermore, pinpointing significant genes within critical pathways offers actionable insights for therapeutic targeting and biomarker discovery. These results underscore PathX-CNN’s capacity to achieve state-of-the-art predictive performance and deliver biologically meaningful interpretations, laying a robust foundation for future experimental validation and application to broader cancer contexts. The present study is based on cohort or class-specific significance. In our future study, we would like to address patient-specific significance to aid personalized treatment strategy.

