## Supplementary File for "PathX-CNN: An Enhanced Explainable Convolutional Neural Network for Survival Prediction and Pathway Analysis in Glioblastoma"

### PathX-CNN Architecture

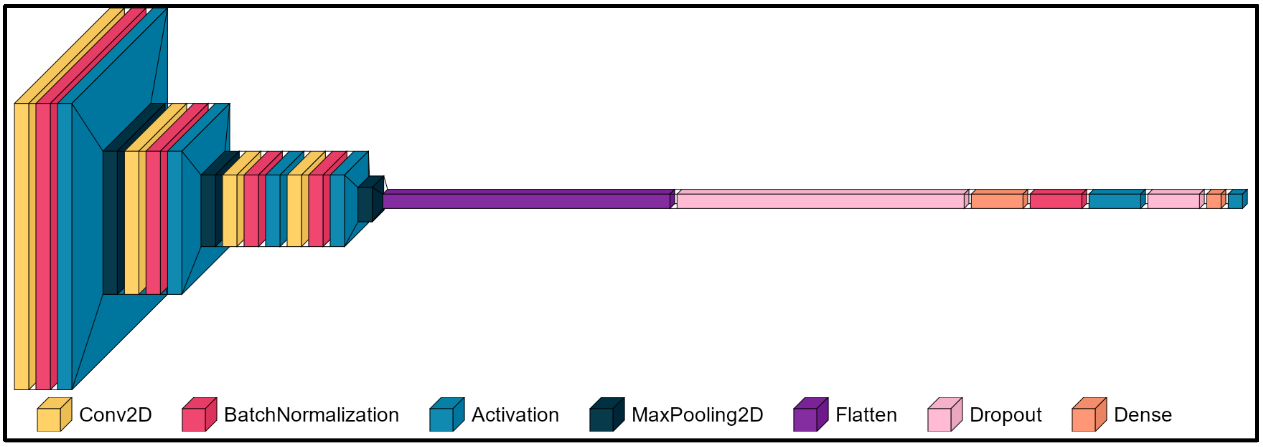


Figure S1. CNN architecture used in PathX-CNN.

### Table S1. Hyperparameter tuning using Bayesian Optimization

| HP | Block1 filters | Block2 filters | Block3 filters | Block4 filters | Dropout 1 | Dense Unit | Dense activation | Dropout 2 | Optimizer | Learning Rate | Decay |
|---|---|---|---|---|---|---|---|---|---|---|---|
| Ranges | 16, 32, 64, 128 | 32, 64, 128, 256 | 64, 128, 256, 512 | 128, 256, 512, 1024 | Min: 0.0, Max: 0.5 step=0.05 | Min: 32 Max: 1024 step=32 | "relu", "tanh", "sigmoid" | Min: 0.0, Max: 0.5 step=0.05 | 'adam', 'rmsprop', 'sgd', 'adamw', 'adadelta' | min: 1e-4 max: 1e-2 sampling:"LOG" | min:0.001 max:0.1 sampling:"LOG" |
| Best HP: Fold 1 | 64 | 16 | 128 | 128 | 0.6 | 720 | tanh | 0 | adam | 0.0003069 | 0.0905127 |
| Best HP: Fold 2 | 16 | 128 | 64 | 128 | 0.6 | 592 | tanh | 0.6 | adam | 0.00013317 | 0.0019706 |
| Best HP: Fold 3 | 16 | 128 | 64 | 128 | 0.6 | 592 | tanh | 0.6 | adam | 0.00013317 | 0.0019706 |
| Best HP: Fold 4 | 16 | 128 | 64 | 128 | 0.6 | 592 | tanh | 0.6 | adam | 0.00013317 | 0.0019706 |

| | | | | | | | | | | |
|---|---|---|---|---|---|---|---|---|---|---|
| **Best HP: Fold 5** | **64** | **16** | **128** | **128** | **0.6** | **720** | **tanh** | **0** | **adam** | **0.0003069** | **0.0905127** |

### Data Description (Details of section 2.10)

The gene-level copy number variation (CNV) was estimated using GISTIC2 [1] method for KIRC, LGG, LUAD and LUSC data. The copy number profile was experimentally measured using whole genome microarray at a TCGA genome characterization center. The TCGA FIREHOSE pipeline then employed GISTIC2 method to generate segmented CNV data, which was subsequently mapped to genes to provide gene-level estimates. The mapping of genes to human genome coordinates was performed using the UCSC Xena HUGO probeMap. Before preprocessing, the CNV datasets for KIRC, LGG, LUAD, and LUSC consists of 528, 513, 516, and 501 samples, respectively. Each of these samples has 24,776 features, representing the copy number variations across the genome.

"MC3" gene-level mutation data for KIRC, LGG, LUAD and LUSC were downloaded from Xenabrowser where the data type is binary (1: non-silent mutation & 0: wild type). Mutation (MUT) data initially comprised of 40,543 features across 368, 511, 513 and 480 samples for KIRC, LGG, LUAD and LUSC, respectively.

In KIRC, LUAD, and LUSC, the DNA methylation (MET) data was sourced from two platforms: Human Methylation 27k (27k) and Human Methylation 450k (450k). LGG has only Human Methylation 450k (450k) data. Initially, the KIRC, LUAD, and LUSC methylation data consists of 418, 151, and 160 samples with 27,578 features and 480, 492, and 415 samples with 485,577 features respectively. Intermediate processing, combining both datasets by retaining the common probes, resulted in 892, 631 and 575 samples with 25,978 features in KIRC, LUAD, and LUSC cancer respectively. LGG contains 530 samples with 485,577 features from Human Methylation 450k platform.

The KIRC, LGG, LUAD and LUSC datasets consist of 606, 530, 576 and 553 samples, respectively, with 20,530 mRNA expression (EXP) features.

**Processed Data:** The processed datasets, common between multi-omics data, for KIRC, LGG, LUAD and LUSC consist of 360, 508, 498 and 473 samples, respectively, with 10,591 features. For patients' survival classification, long-term survival (LTS) was defined as survival > 3 years after diagnosis, while non-LTS was defined as survival of ≤ 3 years. Individuals whose last follow-up was ≤ 3 years were excluded from further analysis. Finally, the LTS group consists of 188, 153, 121, and 134 cases, while the non-LTS group had 61, 75, 123 and 140 cases for KIRC, LGG, LUAD and LUSC, respectively.

**Confusion Matrix from Different Classifiers**

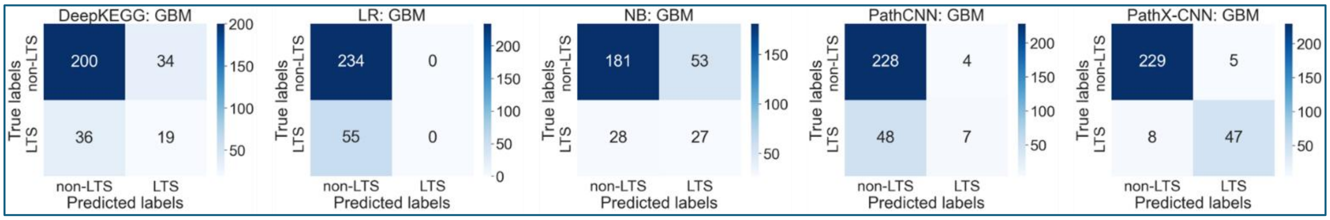


Figure S2. Confusion matrix of GBM data by five different classifiers. LR: logistic regression, NB: naïve Bayesian classifier

**Global Interpretation of Features:**

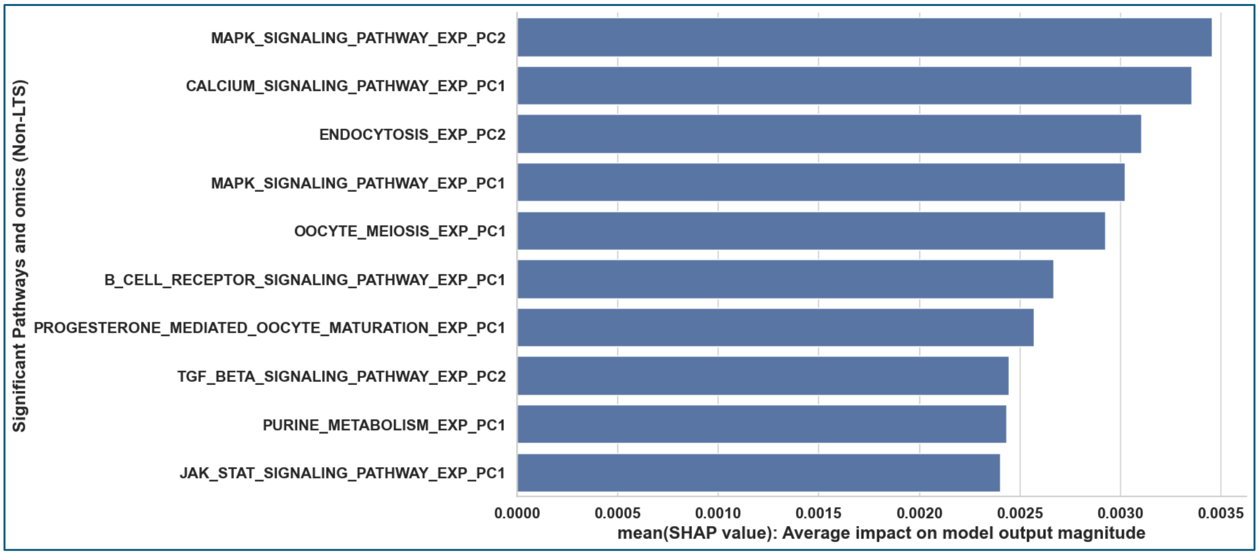


Figure S3. Global Interpretation of Features (PC values of Pathways and omics) using SHAP. This figure shows only the top 10 significant global features for non-LTS class of GBM determined by SHAP score.

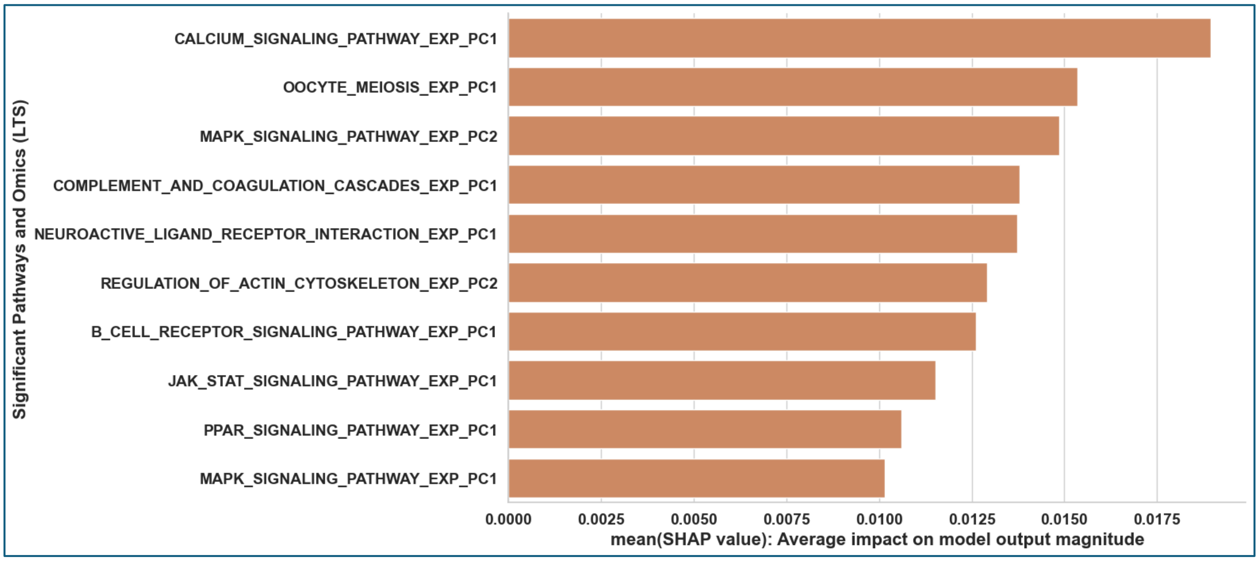


Figure S4. Global Interpretation of Features (PC values of Pathways and omics) using SHAP. This figure shows only the top 10 significant global features for LTS class of GBM determined by SHAP score.

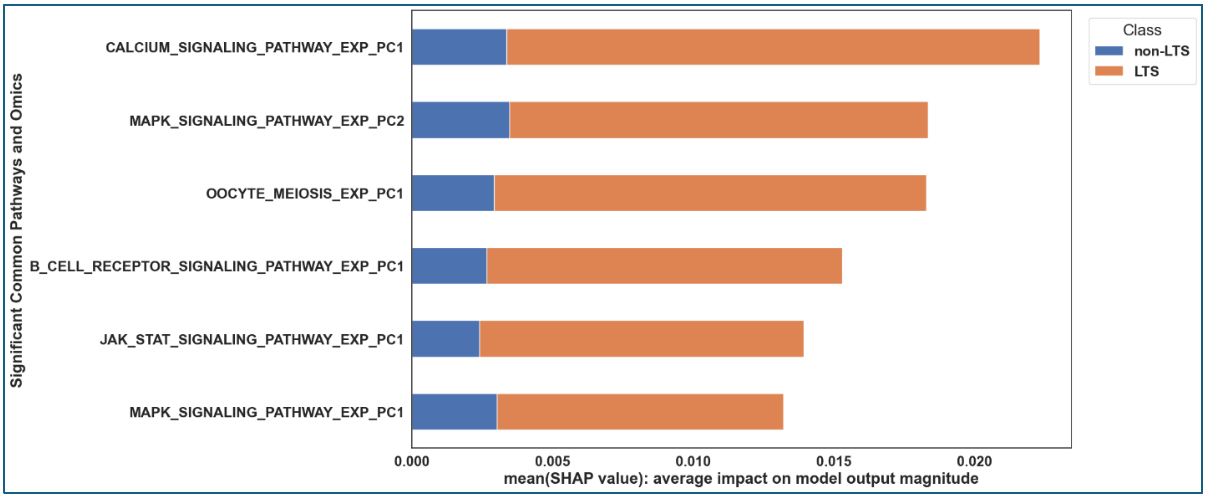


Figure S5. Global Interpretation of Features (PC values of Pathways and omics) using SHAP. This figure shows only the common global features among the top 10 features from LTS and non-LTS class of GBM determined by SHAP scores.

**Significant Genes in the Significant Pathways (Section 3.6)**

| Common Pathways | Significant Genes | # Significant Genes | Total # genes in the pathway |
|---|---|---|---|
| B cell receptor signaling | AKT2, NFKB1, NRAS | 3 | 75 |
| Calcium signaling | ATP2B1, CALM2, GRIN2D, HTR2B, ITPKA, PHKG1, PLCB2, TACR2 | 8 | 178 |
| JAK STAT signaling | LEP | 1 | 155 |
| MAPK signaling | AKT2, AKT3, ATF2, CACNA2D1, CACNG4, CHUK, DDIT3, DUSP1, DUSP14, DUSP5, FAS, FASLG, FGF1, FGF18, FGFR1, GADD45B, GADD45G, GNG12, IL1R1, JUN, MAP2K2, MAP2K5, MAP3K3, MAP3K7, MAP4K2, MAPK13, MAPK14, MAPK3, MAPK9, MAX, NLK, NRAS, NTRK2, PAK1, PAK2, PDGFB, PLA2G3, PPM1B, PPP5C, PTPRR, RELA, TAOK2, TGFB2, TGFBR2, TRAF6 | 45 | 267 |
| Oocyte meiosis | ANAPC10, IGF1, IGF1R, MAPK3, PPP2R1B, PPP2R5C, PRKACB | 7 | 113 |

Figure S6: This figure represents a tabular format to show significant genes in common pathways across LTS and Non-LTS class of GBM. The significant genes were identified using the CoxPH model, with a significance threshold of p-value < 0.01.

Figure S6 presents the significant genes identified in biological pathways common to both Long-Term Survival (LTS) and Non-Long-Term Survival (non-LTS) groups of Glioblastoma Multiforme (GBM). The analysis was performed using the Cox Proportional Hazards (CoxPH) model. To determine significance, a threshold of p-value < 0.01 was applied. The table is structured into four columns: (1) Common Pathways, which lists pathways shared between LTS and non-LTS groups, (2) Significant Genes, showing the genes identified as significant predictors of survival within each pathway, (3) # Significant Genes, representing the count of statistically significant genes in each pathway, and (4) Total # Genes in the Pathway, which provides the total number of genes annotated to the respective pathway. The MAPK signaling pathway contains the highest number of significant genes, followed by the Calcium signaling pathway, Oocyte meiosis, B cell receptor signaling, and JAK-STAT signaling pathways, with 45, 8, 7, 3, and 1 significant gene, respectively.

**Forest Plot for these Significant genes with Hazard ratio and 95% Confidence Interval is shown in following figures:**

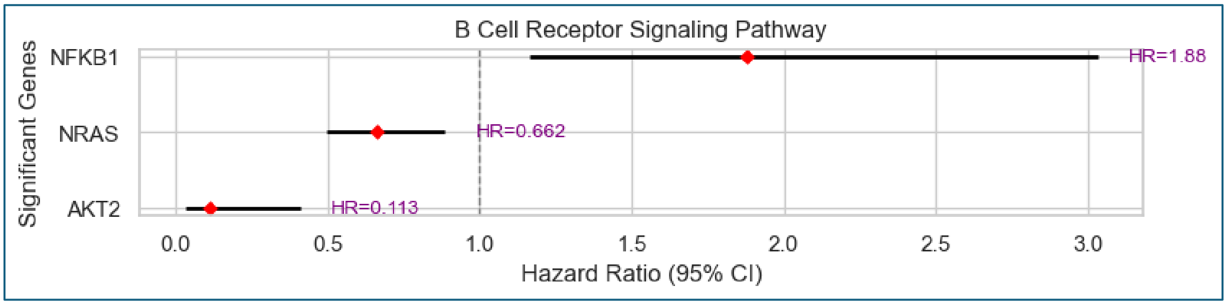


Figure S7: Forest plot of significant genes in B Cell Receptor Signaling Pathway

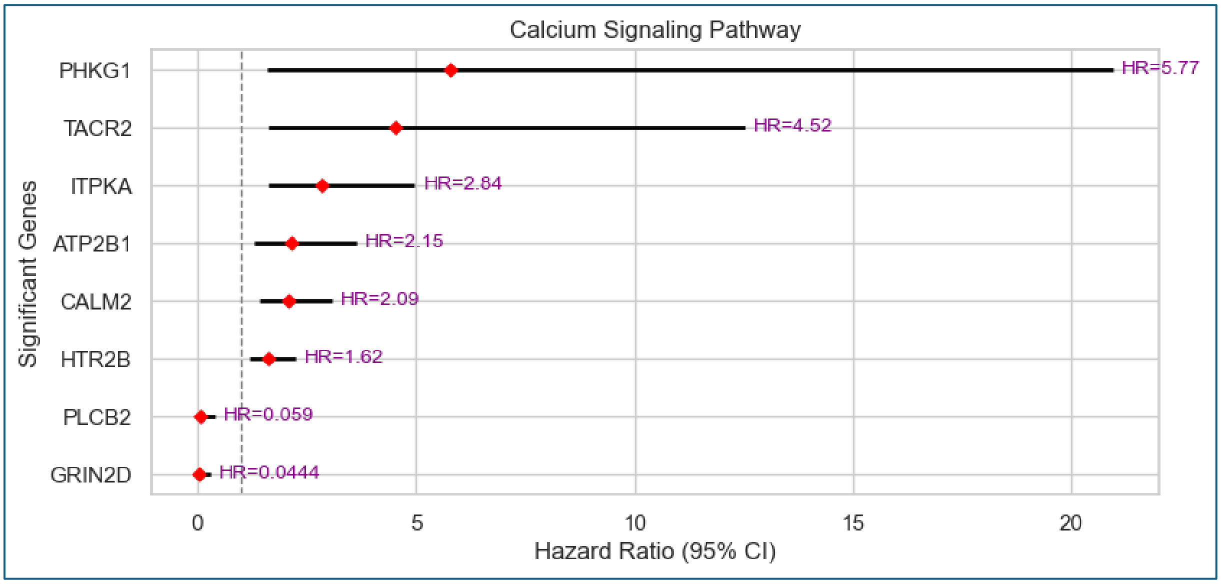


Figure S8: Forest plot of significant genes in B Calcium Signaling Pathway

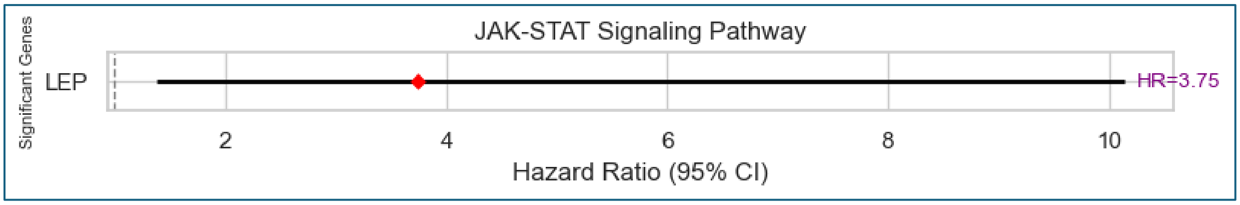


Figure S9: Forest plot of significant genes in JAK STAT Signaling Pathway

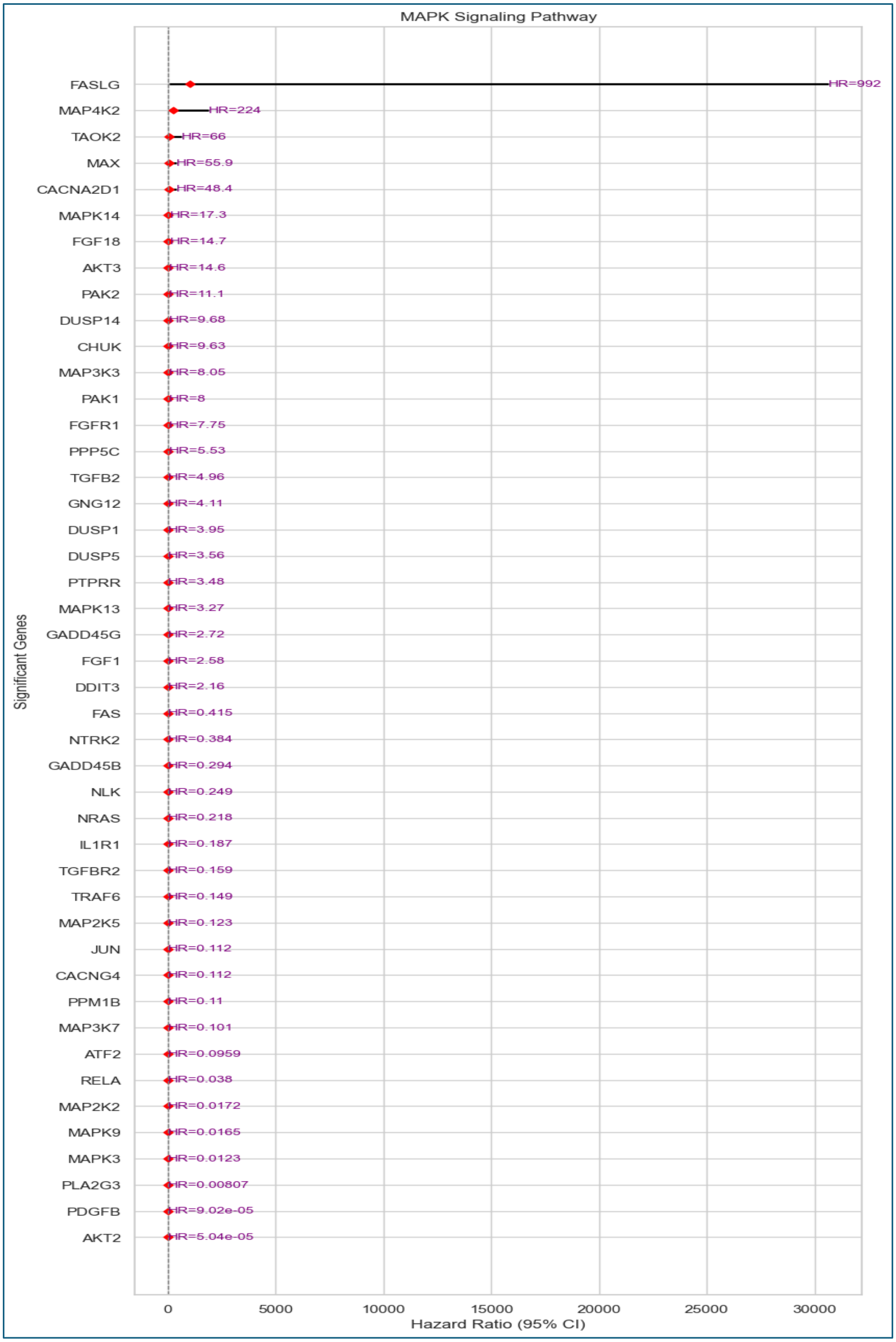


Figure S10: Forest plot of significant genes in MAPK Signaling Pathway

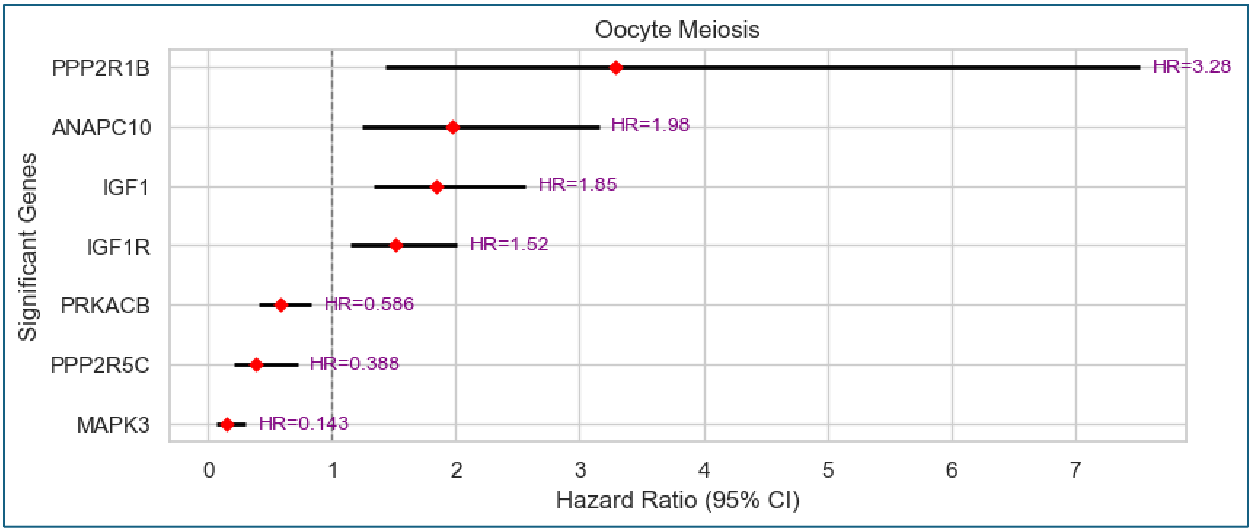


Figure S11: Forest plot of significant genes in Oocyte Meiosis
